# In-depth analysis of proteomic and genomic fluctuations during the time course of human embryonic stem cells directed differentiation into beta cells

**DOI:** 10.1101/2020.10.05.326991

**Authors:** Bogdan Budnik, Juerg Straubhaar, John Neveu, Dmitry Shvartsman

## Abstract

Pluripotent stem cells (PSC) endocrine differentiation at a large scale allows sampling of transcriptome and proteome with phosphoproteome (proteoform) at specific time points. We describe the dynamic time course of changes in cells undergoing directed beta-cell differentiation and show target proteins or previously unknown phosphorylation of critical proteins in pancreas development, NKX6-1, and Chromogranin A (CHGA). We describe fluctuations in the correlation between gene expression, protein abundance, and phosphorylation, which follow differentiation protocol perturbations of cell fates at all stages to identify proteoform profiles. Our computational modeling recognizes outliers on a phenomic landscape of endocrine differentiation, and we outline several new biological pathways involved. We have validated our proteomic data by analyzing two independent single-cell RNA sequencing datasets for in-vitro pancreatic islet productions using the same cell starting material and differentiation protocol and corroborating our findings for several proteins suggest as targets for future research.

Moreover, our single-cell analysis combined with proteoform data places new protein targets within the specific time point and at the specific pancreatic lineage of differentiating stem cells. We also suggest that non-correlating proteins abundances or new phosphorylation motifs of NKX6.1 and CHGA point to new signaling pathways that may play an essential role in beta-cell development. We present our findings for the research community’s use to improve endocrine differentiation protocols and developmental studies.

## Introduction

Human pluripotent stem cells show significant promise for the development of tissuespecific cell types that are used in large-scale screening assays, clinical trials and studies in regenerative biology. The complexity of signaling involved in directed stem cell differentiation varies significantly between cell lineages, but one of the most thoroughly examined cases is that of insulin-secreting beta-cells. Physiological disorders that result in dysfunction of the beta cell, following an immune attack and destruction in Type 1 diabetes (1) or cellular dysfunction as occurs in Type 2 diabetes (2). A promising approach for the beta cell replacement has emerged by the large-scale production of endocrine lineage cells and beta cells from stem cells (3–7).

Directed stem cell differentiation is an array of signaling pathways, controlled with a temporal precision, aiming to mimic development sequences during normal embryonic development. Beta-cell in vitro differentiation spans more than thirty days, with the addition of five different media formulations containing twelve growth factors and small chemicals that regulate ten different cell signaling pathways. Cells progress gradually through distinct phases of endoderm lineage development, resulting in the appearance of beta and other pancreatic lineage cells (5,8). The protocol produces 20-40 % insulin-secreting beta cells, leaving open the ongoing challenge to improve its efficiency.

Transcriptomics studies for human stem cell differentiation for a number of lineages have been performed using bulk RNA sequencing (9–13) and single-cell RNA sequencing (14–16), revealing numerous temporal profiles for transcripts associated with beta-cell development, but similar studies of proteins are lagging. Technological developments in mass spectrometry and nano-liquid separation chromatography (nano-LC) allow us to examine proteomes more extensively and several such studies were reported in recent years (17–19). The in vitro differentiation protocols make it possible to examine both proteome and RNAseq data in the same cell populations at the different stages of the differentiation time course, while providing an ample sample amount to be used for all assays from a single differentiation run. When compared to protein and transcripts—changes, including a number of genes involved, all processes have a slightly reduced correlation of either up-or downregulation for each gene, owning to post-translational modifications at protein levels such as phosphorylation and transcriptome modifications (17,20,21).

Using an extensive transcriptomic, proteomic and phosphoproteomic analyses at each stage of the differentiation protocol, we identified more than 12,000 transcripts and proteins (in each dataset), combined with 4,000 phosphorylated counterparts, all providing a broad view of proteogenomic (phenomic) networks and signaling events that drive the differentiation of embryonic stem cells (ESCs) into endocrine lineage cells in vitro. Our analysis identified proteins with dynamics of mRNA transcripts, where equivalent temporal profile of expression does not follow the same temporal profiles. Computational models of fluctuations in levels of transcripts and their proteins, combined with signaling interpretation of posphoproteomics provided insights in to pathways driving beta cell differentiation. These pathways offer new targets for perturbation via inhibition or activation, as an additional step for translational development and therapeutic implementation of replacement beta-cell therapies.

In order to associate protein targets of interest from the proteome and phosphoproteome datasets with a specific timepoint and specific cell population within the differentiating clusters, we have used unbiased transcriptional profiling and used a visualization method of t-Distributed Stochastic Neighbor Embedding (t-SNE) which allows a non-linear dimensionality reduction for large and complicated datasets such as single-cell RNA sequences (sc-RNAseq) (22). The t-SNE method was applied to two independent datasets for embryonic stem (ES) cell differentiation, using same differentiation protocol and ES starting line (14,23). We show that several highly-expressed or highly-phosphorylated proteins identified by us at later stages of the differentiation protocol can also be attributed to a specific endocrine lineage and may present an interesting target for a future research.

An unexpected discovery during the phosphorylation analysis has occurred, where we have identified three novel and active phosphorylation motifs for key protein driver of endocrine differentiation, NKX6.1, which have not been reported previously. This exciting find accompanies the discovery of post-translational phosphorylations for other proteins, such as Chromogranin A (CHGA, highly phosphorylated at late stages of the protocol), Glucagon, PRRC2A and others involved in beta-cell function and diabetes onset. Our results can be explored further by developmental and translational researchers, studying pancreatic development and cell lineage with its islets.

## Results

### Generation and analysis of transcriptomic and proteomic data of beta-cell differentiation in vitro

To gain insights into the complexity of molecular signaling events during beta-cell differentiation protocol (schematic of pathways involved, Fig. 1A) we collected four independent samples for each stage of the protocol, including the undifferentiated HUES8 ESC clusters, as shown in Fig.1B. It resulted in 4×6 matrices of RNAseq, proteomics and three sets of phosphoproteome raw data, processed as described in Methods. The efficiency of the betacell differentiation protocol was monitored using the expression parameters of endocrine lineage stage markers (3) and quantified by FACS (example FACS plots in Supp. Fig. 1A): SOX17 for definitive endoderm stage (DE stage, mean of positive population ~87%, 20 differentiations including the 4 sets for the analysis here), PDX-1 (~90%) for pancreatic-specific endoderm (PP1), NKX6.1/PDX-1 (~40%) for pancreatic-bud stage (PP2) and NKX6.1/C-peptide for beta-cell formation stage (~14% at day 1, EN and detected via IHC in clusters (Supp. Fig. 1B). At the EN stage, other pancreatic endocrine cells are present, as shown by the expression of GLP2 (α-cells) and Somatostatin (δ-cells).

**Figure 1.**
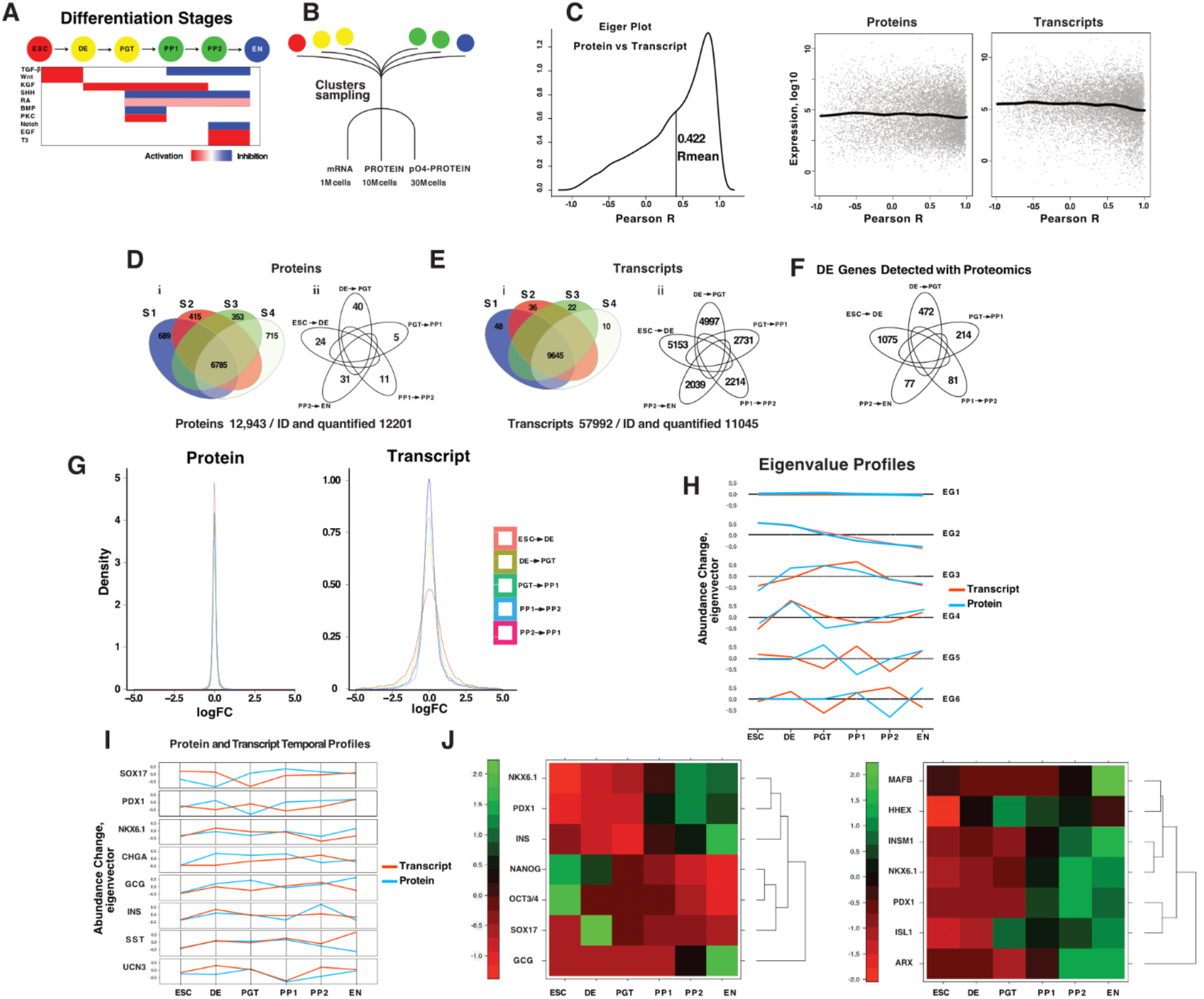
Proteomics vs. transcriptomics analysis and population dynamics during stem cell differentiation towards endocrine lineages. **A.** Description of beta-cell differentiation protocol with key pathways activated (red) or inhibited (blue) as ESCs transition through stages of it. **B.** Sample collection diagram. Cell clusters were collected at each protocol stage and cell count was performed. **C.** Correlation “Eiger” plot of transcript vs. protein levels of expression throughout the differentiation protocol. **D.** Venn diagrams of proteomics sample data distributions (i) between 4 independent samples (S1-S4) with the amount of UNIPROT-identified proteins and (ii) amounts of differentially expressed proteins (p<0.05) between each stage of beta-cell differentiation in vitro. The full protein name, fold change list is shown in Supp. Table 1. **E.** Venn diagrams of RNAseq sample data distributions (i) between 4 independent samples (S1-S4) with the amount of UNIPROT-identified mRNA transcripts and (ii) amounts of differentially expressed unique mRNAs (p<0.05) between each stage of beta-cell differentiation in vitro. **F.** Venn diagram of differentially-expressed (p<0.05) transcripts between each stage of the beta-cell protocol, identified both in RNAseq and proteomics data analysis. DE-definitive endoderm, PGT-primitive gut tube, PP1-progenitor pancreas stage 1, PP2-progenitor pancreas stage 2, EN-endoderm. **G.** Density plot of total protein expression values at each protocol stage sample vs. log (10) fold-change values of protein expression between the stages. Density plot of total mRNA expression values at each protocol stage sample vs. log (10) fold-change values of mRNA expression between the stages. **H.** Eigenvector profiles dynamic clustering for protein and mRNA expression profiles. Derivation from 0.0 value indicates an observed change in RNA/protein expression. **I.** Beta-cell protocol stage markers Eigenvector dynamic profiles for protein and mRNA expression during the differentiation transition. **J.** Heat-map for the expression of individual stage markers during beta-cell protocol (p<0.05), color coded for t-statistics range (−1.75,2.25) and clustered.

RNAseq data of 11045 (from 57992 of total mRNAs detected) unique transcripts were identified with Uniprot and TREMBL proteomic databases and used to generate a list of mRNAs cross-referenced with proteomic nano-LC data. Proteomic datasets were made using standard nano-LC protocols from the same replicates as RNAseq datasets, with 12,201 unique protein IDs identified and quantified from total of 12,943 proteins to be identified.

First, we have created “Eiger” plot (named after a characteristic profile, resembling Eiger mountain in Switzerland) of Pearson correlation coefficients R for same gene IDs identified in each dataset, between proteome and transcriptome (black line, R protein). The mean R value for proteome/transcriptome correlation was 0.422, indicating a partial or weak correlation between two datasets (Fig.1C). The distribution of protein and transcript populations between the obtained R values is shown in Fig.1C. We show that described populations are distributed approximately uniformly between all R values, and there was no significant shift in the distribution, regardless of R value and either proteome or transcriptome datasets, as indicated by no marked change in fitting curves (blue line). (Fig.1C).

Venn diagrams were generated for proteomes (Fig.1D) and transcriptomes (Fig.1E) and differentially expressed proteins (see the full list in Supp. Table 1) were identified by statistical evaluation of the changes in their amounts (p<0.05) between each differentiation stage. The transcriptomic dataset as generated by the current methods have a significantly higher coefficient of variance than the proteomic counterpart (Supp. Fig.2), which can be attributed either to some methodological differences or the intrinsic variability between replicates at transcript levels. As shown in Fig. 1E, the number of differentially expressed (DE) mRNAs were also several orders of magnitude higher than the amounts of differentially expressed proteins. To understand further the correlation between differentially expressed transcripts and differentially expressed proteins, we generated Venn diagram only with mRNA that can be cross-referenced with protein IDs. The number of differentially expressed mRNAs was several orders of magnitude higher than proteins (Fig.1F). The magnitude of the change in transcript levels and numbers of transcripts compared to their protein counterparts is shown in Fig.1G. Overall protein amounts (density in Fig.1G) and the changes in those levels showed less volatility than transcriptome values at each stage of the beta-cell protocol. This analysis points to the conclusion that significant changes at the transcriptome level do not always translate to the same shift at protein levels from the same gene or gene groups.

To outline the dynamics of transcriptome and proteome of expression during beta-cell differentiation, we generated and clustered (see Methods) dynamical expression profiles (Eigenvectors) for transcriptome and proteome (24) via unsupervised machine learning (25,26). Each Eigenvector profile represents the change in dynamics (“vector direction”) of a certain gene or gene cluster as it proceeds through the protocol stages. Our analysis resulted in 6 groups of mRNA/Protein Eigenvalue patterns (Fig.1H). The x-axis values represent the observed change in gene expression, where positive or negative values indicate only the observed Eigenvector value and are not the increase or the decrease in levels of a specific transcript or its protein. Two of the six identified clusters do not follow the expected change in protein expression upon the change in a transcript, and there are temporal shifts or even delays in protein expression pattern (Fig. 1I, clusters EG5 and EG6).

We also generated individual Eigenvalue profiles for endocrine lineage marker proteins, SOX17 for definitive endoderm, PDX1 for pancreas specific-endoderm cells, NKX6.1 for pancreatic progenitors, INS for a beta, GCG for alpha, SST for delta cells and UCN3 These profiles show (Fig.1I) that the observed and the expected changes in transcript profiles do not always correlate with the change in the protein. This observation emphasizes the fluid behavior of protein expression lineage cell markers, underlining the importance of their identification at each stage both at protein and transcript levels. To validate the expression of the expected markers at each stage of the beta-cell differentiation protocol, we generated heat-maps from proteomics data, with the exception for UCN3 which was not detected in all three proteome sets (Fig.1J).

### Identification of proteins within the signaling pathways involved in beta-cell differentiation protocol

To confirm the presence of protein targets perturbed by the differentiation protocol, we profiled each signaling pathway protein component either activated or inhibited by the protocol signaling proteins or compounds (Fig.2 and Supp. Fig.3). We show the distribution of KEGG identified signaling pathways and Pearson correlations (R’s) between proteome and transcriptome datasets analysis (Fig.2A). We set the cut-off for “good correlation” at R > 0.6 and for the “poor correlation” R < 0.5. All “good” correlation proteins were analyzed by KEGG pathway analysis, same for the “poor” ones. We have found 125 KEGG pathways with “good” R correlation proteins, 32 with “poor” and 54 KEGG pathways had proteins from both “good” and “poor” proteome datasets. Using independently developed method for identifying distinct signaling pathways clusters/programs during beta-cell differentiation, using multikernel learning (SIMLR) and topic modeling analysis of single-cell transcriptomic data (23), we show that both proteome and transcriptome, in general, conform to a linear relationship between stages of the protocol (Fig. 2B).

**Figure 2.**
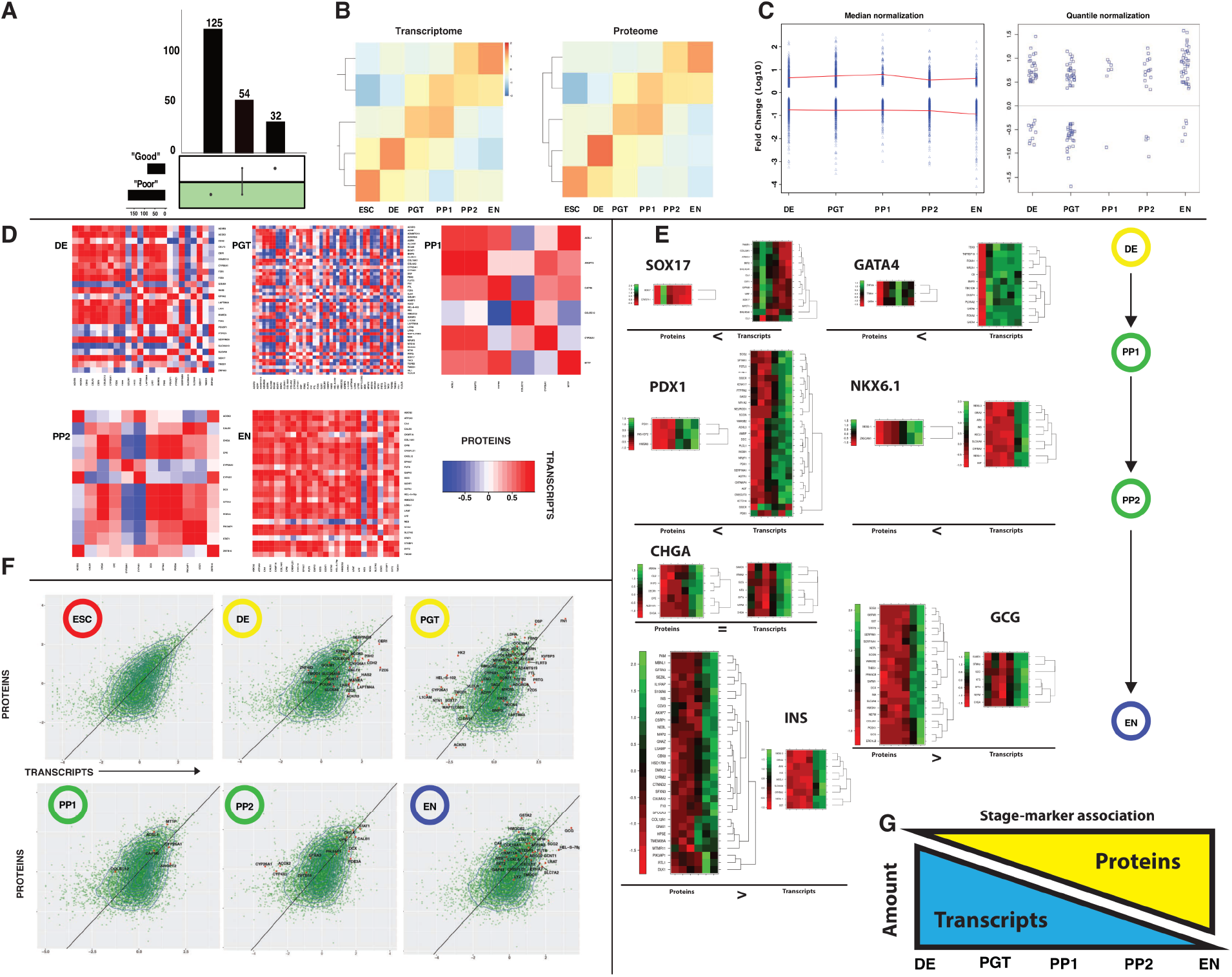
Proteomic examination of signaling pathways induced by beta-cell differentiation protocol. **A.** Proteome data were split in two groups (“Good” Correlation R>0.5 and “Poor” Correlation R<0.5), after which KEGG pathway analysis was done. All identified pathways were cross-referenced and Venn diagram was generated. **B**. Transcriptome and proteome heatmaps thorough the stages of beta-cell differentiation protocol, each row represents same gene ID in both datasets. **C.** Scatter plots of Log (10) fold-change (logFC) of differentially expressed DE proteins, after median and quantile normalizations, plotted against each stage of beta-cell differentiation protocol. **D.** Correlations of DE proteins and corresponding mRNA transcripts at each stage of the beta-cell differentiation. **E.** Unsupervised clustering of proteins and transcripts with stage-specific endocrine markers. **F.** Landscape of total transcripts and proteins through stages of differentiation with annotated DE protein IDs. **G.** Correlation diagram for the number of transcripts or proteins clustered with each stage marker as function of the differentiation protocol stage.

### Differentially expressed proteins indicate targets for perturbation that are involved in beta-cell differentiation protocol in vitro

After analyzing proteins and transcripts that were significantly up/down regulated between each stage checkpoint of the protocol, we performed a stage clustering of proteins with different normalization methods (median and quantile)(27,28), plotting their Log (10)-fold change values against the stage checkpoint (Fig.2C). We also generated fitting curves for the total transcriptome fluctuation dynamics (Supp.Fig.4, left panel) and plotted a fold-based plot of protein changes (Supp.Fig.4, right panel). We observe a constant decrease in the number of changing transcripts as cell culture progresses through the protocol stages, although the overall numbers remain several orders of magnitude higher than the proteins at each stage checkpoint of the protocol.

An initial method based on a simple comparison between protein IDs that are up or down-regulated between two individual protocol stages yields 1000’s of proteins. This large amount of protein IDs makes difficult to researchers to reach a definitive conclusion when dealing with large and dynamic datasets, unless they are looking for a particular gene or family of proteins in it.

To gain some clarity into a complex and dynamic proteome landscape we employed two methods of dataset normalizations: in first, data samples were first normalized individually by centering around the median over all channels. Additionally, a variance stabilizing data transformation (29) was applied to the complete data set and batch effects were removed by regressing on the TMT runs (30). After that, log10 fold-changes of median-normalized protein IDs were plotted at each protocol stage and STRING analysis was done on up/down regulated IDs with a log10 value above or below 0.5 (3-fold change, Supp. Fig. 5) for PP1 to PP2 and PP2 to EN stage transitions (Supp. Fig. 5). We were able to identify several interesting STRING nodes, including APOE node which is up-regulated during the PP1 to PP2 transition and down-regulated during PP2 to EN. Interestingly, the role of APOE in pancreas function has been established by epidemiological surveys and offered some insight in APOE-induced toxicity for the islets (31) and was also identified by single-cell sequencing study as some of the highly changing transcripts (14). We suggest that perhaps APOE function and expression pattern deserve more attention during the beta cell development in vitro. A cluster of AGRN (Agrin) and corresponding integrins (ITGA) appears during PP2 to EN stage, and while the role of integrins in beta cell development and function was widely recognized (32–34), we could not find any evidence of previous studies for the role of Agrin in beta cell life cycle. Lastly, an interesting cluster of kinesin proteins appears in upregulated STRING network of PP2 to EN transition. Kinesins are important for beta cell priming for secretion and creation of the initial population of insulin-containing dense core vesicles docked to plasma membrane (35,36). We interpret this observation as the first sign of endocrine cells entering a maturation (secretion) phase of development.

Another, more rigorous approach to proteomic data normalization utilizes quantile normalization (27,28,37). This type of normalization is used to equalize different amounts of samples injected into the instrument between different samples. For TMT type of analysis, such normalization is used for different channels in the same TMT labeled study; it assumes that the whole proteome among samples is equal, and it normalizes sample load between channels before significantly differential proteins can be revealed. When a biological system undergoes dramatic changes, in such cases, the total proteome is changing too. For such a case, the use of quantile normalization can over normalize proteins changes to the point that only proteins that had been changed dramatically, such as those that are newly produced and been specifically degraded, will show up as differentially expressed after quantile type of normalization. Such a dynamic system assumption of total proteome stability cannot be used, and only mild normalization should be performed to prevent such losses.

The result of differentially expressed proteins at each stage transition, after quantile normalization, (shown in Fig.2C, right panel) shows an unbalanced distribution between the up/down regulated differentially expressed proteins as cells proceed within the protocol, and were selected by having a statistically significant increase in abundance. While at the DE and PGT stages there is an almost equal number of up/down regulated proteins, at the PP1 stage onwards there are more differentially expressed proteins that are upregulated (see Supp. Table. 1). Next, we proceeded to plot each of the differentially expressed proteins with their transcripts (Fig.2D heat-maps in red and blue), showing that while the individual protein and its transcripts highly correlate in the observed change of expression (red color), there are notable differences between the different proteins in the strength of the correlation when compared to each other (blue color). If to compare the amount of correlation noise (red to blue color amount in the heat-map) for protocol stages, we see that from the DE to PP2 stages there is a significant discrepancy between the protein/transcript changes of each differentially expressed protein, and only upon entering the EN stage almost all differentially expressed proteins are synchronized (heat map almost red in color).

To further our understanding of additional signaling pathways in the beta-cell protocol, we set to generate heatmaps for the lineage specific genes and to correlate via unsupervised clustering methods their transcripts and proteins with the RNAseq and proteomic datasets, producing unlimited number of protein and transcript targets in the final heat-map plot. We used endocrine lineage specific stage markers SOX17, GATA4, NKX6.1, PDX1, CHGA, and Insulin (INS) as anchors for the generation of heat-maps as shown in Fig.2E. The overview of both RNAseq and proteomic heatmaps demonstrates unequal distributions between the number of transcripts and proteins that cluster with each marker, with a notable exception of CHGA (full list of clustered transcripts and proteins is shown in Supp. Table 2). As cells progress through stages of the protocol, the amount of proteins clustered with a specific marker gradually increases, while the number of transcripts goes down (see the model in Fig.2G).

Finally, we mapped each differentially expressed protein/transcript reference point (red dots) on the scatterplot for the global landscape of gene expression at each stage of the protocol (Fig.2F, bottom row) in order to see whether protein position within the plot will allow us to identify clusters of differentially expressed proteins, which may indicate common biological processes that involve these targets. We also added the R correlation fit line (black) in each plot serving as the reference for the closely correlated proteins. Our analysis shows that every differentially expressed protein remains at individual positions within the scatterplot and not in clusters. It suggests that each differentially expressed protein can serve as node marker of a particular biological process and may be explored for perturbation for the improvement of beta-cell differentiation in vitro.

### New insights in stage marker-correlated proteins and description of proteome accompanying morphological changes in cellular clusters

Part of production quality controls and assurance (QC/QA) during the beta-cell differentiation in vitro involves continuous monitoring of the cluster morphology, where its size, shape and cell density continually changes upon the progression of the protocol (Fig. 3A, phase-contrast images of whole clusters). These changes can be attributed to cell migration and reorganization within the cluster, as cells differentiate, move and coalesce into cell-specific regions within the cluster that in the end resembles islet morphology (see IHC image of whole cluster section in Supp.Fig.1), as also described in (23). It has been previously noted that interactions with connective tissues, integrin expression and extracellular matrix (ECM) protein composition affects beta-cell generation and pancreatic islet function (14,38–44). We set to correlate one key ECM component isoform (Laminins, LAM) and the ECM binding proteins (Integrins, ITG) detected via proteomics and clustered in heatmaps according to stages of differentiation protocol (Fig.3B). We were able to show that indeed specific isoforms of integrins identified with endocrine populations appear at same the moment when the lineage begins to emerge (PP2 stage, α1 and β1 integrins, ITGA1 and ITGB1 respectively) and their ECM counterpart laminin-511 (subunits LAMA5, LAMC1). Noticeably, laminin-511 subunit LAMB1 appears earlier at the PGT stage, and the same observation stands for the laminin-521 appearance in clusters. In addition, we clustered collagen (COL) isoforms and ADAMTS family of multidomain extracellular proteases, since both are integral for ECM formation and processing, as cells move around the cluster (Supp. Fig.6). Several targets in both clusters were identified to be correlated with each lineage marker, for example ADAMTS16, COL12A1 and COL26A1 with insulin (INS).

**Figure 3.**
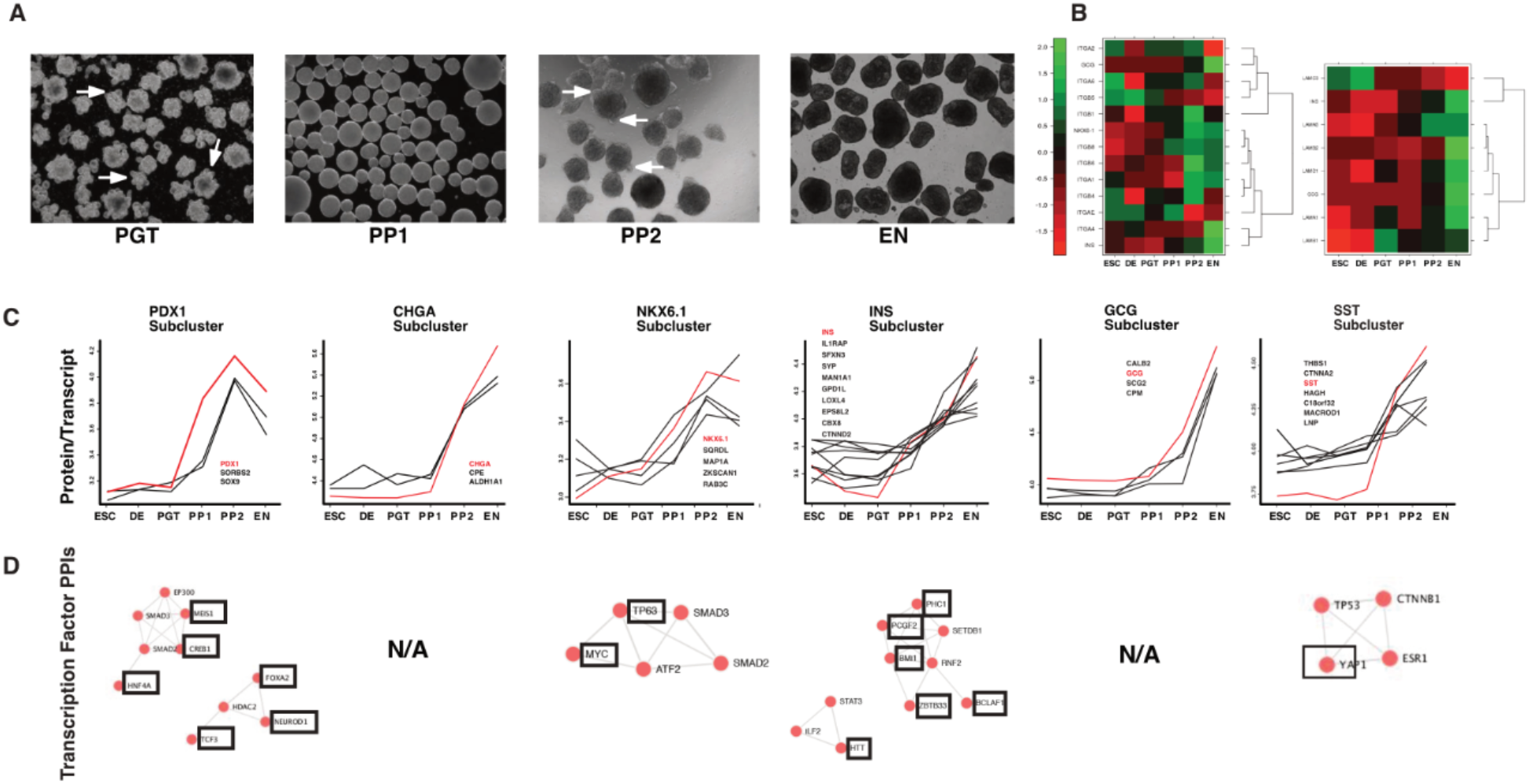
Biological insights from transcriptome and proteome comparisons. **A.** Phase-contrast images (4x) of the morphological change in cellular clusters in suspension as they progress through the stages of the differentiation protocol. **B**. Heat-maps of integrins and laminins, identified at each stage of differentiation. **C.** Protein to transcript ratio curves of stage markers and proteins identified via unsupervised learning. Transcription factor networks generated by ENRCHR analysis of stage marker protein cluster. Statistically significant candidates (p<0.05) are shown in black squares.

ECM collagen and laminin components, its mechanical properties, and integrins interaction with it govern lineage determination of progenitor cells within the developmental niche (45,46), where different laminin and integrin bindings take a significant part, one can use isoforms of these proteins as accessible targets for perturbations. It would enable an enrichment of a target endocrine cell populations present within the clusters.

### New signaling pathways via network analysis and growth factors identified through proteomics

To identify new signaling networks we use machine learning and stage marker proteins as anchor reference genes for generating gene expression profiles that are similar to the anchor (red line in graphs, Fig. 3C). Lists of proteins that were showing similar profiles of those for stage markers via unsupervised clustering loop analysis were generated, where both transcripts and protein abundances were used for the machine learning process. After identifying the targets, ENRCHR tool was used to find transcription factors that interact with these protein IDs in each cluster. Proteins with patterns similar to the specific stage marker can serve as an individual node of biological process, but with the addition of transcription factors detected by the ENRCHR provides another insight into beta-cell protocol signaling (Fig. 3D). Consider the example of the BMI1 protein that was identified through INS cluster analysis and negatively regulates insulin sensitivity and glucose homeostasis (47). YAP signaling factor is implicated in islet development and the increase in beta-cell mass (48–50), but here we detect it only in the somatostatin related network and not in the insulin related network (Fig.3D).

Another attempt to visualize new signaling networks discovered via analysis of differentially expressed transcripts or proteins between each stage of differentiation protocol was done by submitting their respective lists for STRING interaction network analysis (51,52), as shown in Supp. Fig.5. Initial review of transcript-obtained network shows (Supp.Fig.7A), as expected due to a large number of changing transcripts, complex networking arrays between genes, which were hard to interpret, although this complexity gradually diminishes as cells were progressing into the differentiation. Proteinbased STRING network of DE proteins after quantile normalization was smaller in scale (Supp.Fig.7B), with clear nodes of interactions within.

The interpretation of significance for each gene or gene family for endocrine differentiation and identified as a node of interaction with the network has to be taken with a limitation due to the increased heterogenetic population in cell culture. For example, STRING network during the PP1 to PP2 transition shows, cluster of glucose and pyruvate metabolism related genes (DLD, DLAT, DLST, PDHB, PDHX) and ATP-synthesis components (SUCLA2, SUCLG1, SUCLG2) in mitochondria, indicating the change in the metabolic profile of cells. Another example is the NF-kappaB gene (RELA in STRING network) and its related cluster of transcription factors (GTF2F1, GTF2B, GTF2H4, GTF2E2, GTF2E1 and others), indicating possible role of NF-kappaB and its associated signaling for generation of endocrine cells in the protocol, as was stated previously with conflicting conclusions (53,54). STRING network for transcripts during the PP2 to EN transition, shows that cells undergo changes in gene translation (indicated by PCNA and MRPS-family genes), while undergoing another change in metabolic processing of citric acid (IDH3A, IDH3B, and IDH3G), indicative of the possible emergence of insulin secreting beta-cells (55,56).

When STRING analysis was applied to differentially expressing proteins between stages of beta-cell differentiation protocol a variety of clear network nodes have emerged (Supp.Fig.6B): in ESC to DE transition we show WNT pathway cluster, as expected, while identifying new node of fatty acid beta-oxidation (HADHA, HADHB, ACAT1, ACOX3 and others). In DE to PGT transition, beta oxidation node remains, indicating that cells still rely on fatty acid metabolism, while Notch node emerges, combined with ACLY gene of citric metabolism and WNT pathways. In the same transition we show AGR and MUSK genes as nodes of the network, probably correlating with changes in cluster morphology, laminin binding and cell re-organization (57–59). During the emergence of PDX-1+ endocrine progenitors (transition from PGT to PP1) we detected cluster of potassium channel gene family (KCNIP), sodium channel (SCH5A) and DPP6 protein recently reported as beta and alpha cell marker, surprisingly correlated with them, indicating possible new interactions between these proteins (60,61). At the next transition (PP1 to PP2) STAT1 protein emerges as the critical node in the network, correlating with the NF-kappaB network captured by transcript STRING, and with IL-6, Janus kinases (JAK2, TYK2, JAK1) and interferon receptor and its related proteins (IFNAR1, IRF1, IRF9, IFIT1), indicating interferon signaling pathway activity in culture, previously only reported in Type 1 diabetic models of beta cell destruction. Another observation of a signaling network emerges from the STRING network of PP2 to EN transition, where JAK tyrosine kinases, CXCR4 receptors, and STAT pathways are present in culture, while interacting with Ephrin receptor family, as shown previously (62,63) and confirmed here (Supp. Fig. 4B). This network can be an early indication of emerging beta-cell interconnectivity and selforganization within the clusters as also observed by distinct patterns in pancreatic islets (64).

### Phosphoproteomic profiling suggests perturbation targets in beta-cell differentiation

Scalable differentiation protocol used here provides a large sample amount, which can be split between deep sequencing, deep proteome and deep phosphoproteome analyses. Between three datasets we have identified 4275 unique phosphorylated protein IDs (Fig.4A) and have selected 2078 IDs for the initial analysis. Similar “Eiger” plot shows a weak Pearson correlation between levels of phosphoprotein abundances and their transcript counterparts, as generally expected due to two degrees of separation between two gene instances (transcript and phosphorylated protein). Interestingly, PO4-protein R value was not significantly different from the protein R value (Fig1.C, PO4 R-value of 0.375 vs. protein R-value of 0.422). We proceeded to analyze distributions of protein IDs (peptides) with one, two or three PO4 modifications (Fig.4C). Violin plots show a slight increase in the amount of single and double-phosphorylated proteins as cells proceed via the differentiation, while triple-phosphorylation remains a rare event and is decreased from the ESC stage onward.

**Figure 4.**
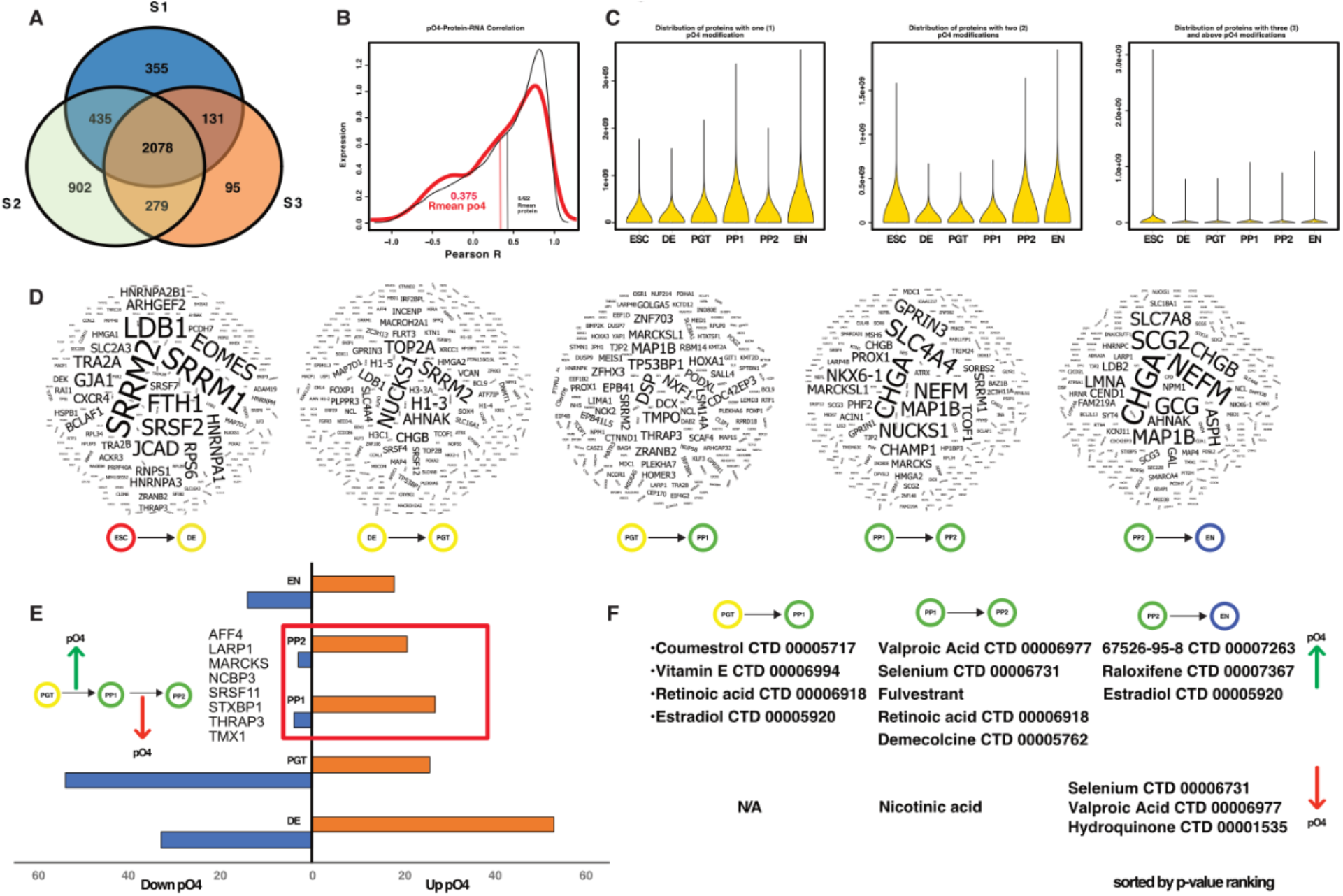
Phosphoproteome analysis of beta-cell differentiation suggests several perturbation targets. **A.** Venn diagram of phosphorylated protein ID distributions between three individual samples datasets. 2078 phosphoprotein IDs were used for all subsequent studies. **B.** Pearson correlation “Eiger” plot of transcript vs. phosphoprotein levels of expression throughout the differentiation protocol. **C.** Violin plots of amount distributions for peptides (protein IDs) with one, two or three phosphorylated amino acid residues. **D.** Word cloud clusters of top 500 phosphorylated peptides (protein IDs), ranked from the highest amount (larger font) to smallest one, at each stage transition during the differentiation protocol. Clusters were generated with online tool (https://www.wordclouds.com/). **E.** Differentially-phosphorylated protein IDs sorted for Up or Down-phosphorylation at each stage of the protocol. Red box outlines stage transitions (PP1 to PP2) where same IDs were down-phosphorylated from being the up-phosphorylated in a previous stage (PGT to PP1). **F.** ERCHR-identified drug compounds for the three last stage transitions in the differentiation protocol, where the phosphoprotein IDs for the drug protein network were either up-or down-regulated.

To identify proteins with highest change in PO4 level during the stage transition and to find novel signaling pathways we have processed top 500 protein IDs at each stage of the protocol and created a “wordclouds”, where highest ID is located in the center and has a largest font size (Fig.4D). We show that during the ESC to DE proteins to have highest phosphorylation are SRRM1, SRRM2, LDB1, FTH1, EOMES, followed by others. For the DE to PGT stage, SRRM2, NUCKS1, TOP2A, AHNAK and H1-3 proteins were at the lead; for PGT to PP1 transition list of leading protein has grown with the notion of DSP, NXF-1, TP53BP1, HOXA1, TMPO and others. For the PP1 to PP2 transition, we identified CHGA as the leading phosphorylated protein, followed by NEFM1, NUCKS1, SLC4A4, GPRIN3, CHAMP1, PROX1, CHGB and, unexpectedly, by NKX6.1 protein which is a key protein for pancreatic development. CHGA maintained the lead as the most abundant phosphoprotein during the final PP2 to EN transition and was trailed by Glucagon (GCG), NEFM, SCG2, CHGB, SLC7A8, MAP1B, AHNAK and ASPH. Beside the already established proteins involved in endoderm development (NKX6.1, CHGA, GCG), we show many other which can serve as anchors or identifiers for a specific biological process or pathway during the differentiation. We have found a limited number of publications showing the phosphorylation of Glucagon or Chromogranin, with no publications demonstrating the phosphorylation of the NKX6.1.

Next, we proceeded to try identifying specific signaling pathways which could be the next target for perturbations for the differentiation protocol improvement. We hypothesized that by clustering of differentially phosphorylated proteins a looking for pathways which are upregulated in one stage and down regulated in the next one (Fig.4E) This type of pathway can be an interesting target for modulation seeking the improvement of the differentiation. We show that stage transitions PGT to PP1, PP1 to PP2 and PP2 to EN have the largest number of protein IDs with an increase in their phosphorylation levels. Our goal was to identify a possible chemical compound or known drug, which can quickly be put into screening assay using beta-cell differentiation protocol

Using the ENRICHR tool we have sorted the results from one of ENRCHR-connected databases DSigDB: Drug Signatures Database, published by (65) and results of the p-value ranked compounds are shown (Fig.4F). We show that pathways associated with the addition of retinoic acid (RA) are detected, which is in correlation with the differentiation protocol first addition of RA from GTE to PP1 and continued with a lower dose from PP1 to PP2. In addition, vitamin E, estradiol and coumestrol compounds were identified. All compounds shown to be important for the function and development of islets in vivo, however it would be interesting to evaluate their effect during the beta-cell production in vitro. At the PP1 to PP2 transition valproic acid emerged as interesting candidate compound, which pathway is upregulated and then downregulated at PP2 to EN. Importance of an essential trace element selenium (Se) in upregulation of beta-cell specific gene expression and insulin content is established and it is used as supplement during the beta-cell differentiation. We also confirm the presence of Se-activated pathway during the PP1 to PP2 transition, which may serve as a point of adding external Se to a defined media. The PP2 to EN transition shows a decrease of Se-related phosphorylation, pointing to a possible exhaustion of external selenium and the need to add more of it into the media. Estradiol and other selective estrogen receptor modulating compounds (fulvestrant, estradiol and raloxifene) pathways are shown to be activated during PP1 to PP2 and PP2 to EN transition. While the estrogen receptor pathway is studied extensively in animal models of pancreas development and diabetes, we propose to use it as a defined target for the activation during later stages of beta-cell differentiation in vitro.

### Identification of new phosphorylation sites for NKX6.1 and Chromogranin A (CHGA)

While both NKX6.1 and Chromogranin A are broadly recognized as key proteins regulating islet growth and markers for the successful differentiation in vitro we could find a little to none evidence for their phosphorylation profiles and role they play in pancreatic development and function. As shown in Fig.5A, a major uptake in phosphorylation levels of the general protein population takes place during the PP2 stage and subsides toward the EN. While DE, PGT and PP1 stages have a large population of detected PO4 peptides, we choose to focus and to highlight those which were reported to be involved in beta cell development and a few with an unexpected high level of phosphorylation. The highest amount of PO4 peptides was detected for CHGA with second spot taken by NEFM (medium neurofilament protein) at PP2 stage. While CHGA appearance is expected in endocrine differentiation, the high levels of NEFM remain to be studied further, although it is detected in several single-cell sequencing studies (66,67). We highlighted few protein IDs that were in top 500 of highly-phosphorylated peptides detected between three datasets (SLC4A4, NKX6.1, PPRC2A, Glucagon (GCG) and Secretogranin II (SCG2). Both CHGA and NKX6.1 presented us with an interesting opportunity to pinpoint each serine or tyrosine phosphorylation sites and to analyze the dynamics of it.

**Figure. 5.**
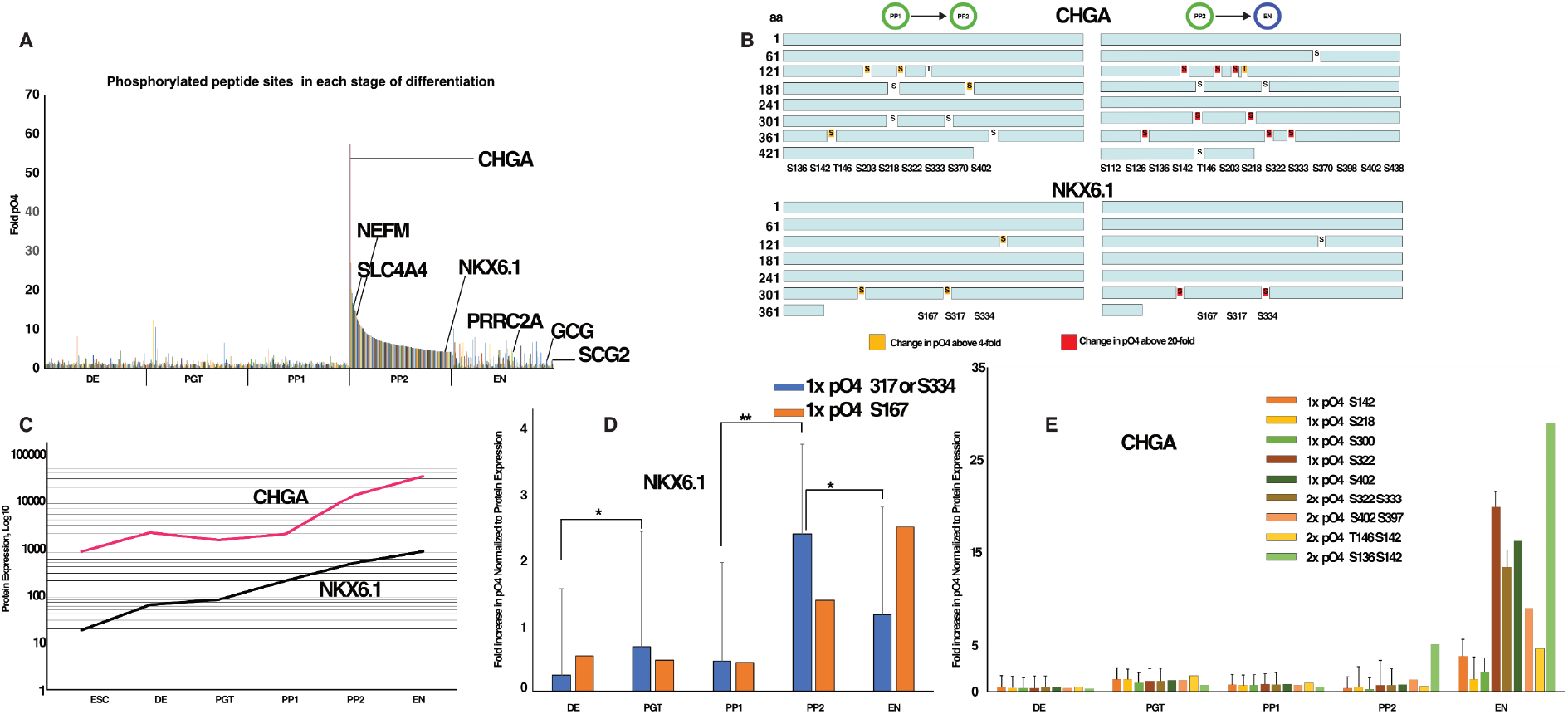
Chromogranin A (CHGA) and NKX6.1 amino acid residue identified as targets for phosphorylation during the PP1 to PP2, and PP2 to EN stage transitions. **A.** Summary graph of fold changes phosphorylation levels for top-500 protein IDs. **B.** Protein sequence for CHGA and NKX6.1 with Serine (S) or Threonine amino acids identified as phosphorylation targets in peptide sequences. **C.** Change of expression (total detected protein amount) for CHGA and NKX6.1 on Log (10) scale for the duration of differentiation protocol. **D.** Changes in phosphorylation of NKX6.1 and CHGA for one or two phosphorylated amino acids in a peptide sequence, normalized to changes in total NKX6.1 and CHGA protein amounts. Bars with error bars have three replicates, whereas no error bars indicated less replicates.

We detected a four-fold increase in CHGA PO4 levels at indicated serine and one threonine sites (Fig5.B, upper panels) during the PP1 to PP2 transition, which was exuberated upon the PP2 to EN transition, with additional serine sites showing phosphorylation. There are several studies describing CHGA phosphorylation (68–70), but we suggest that it deserves a larger attention from the research community, due to observed dynamic phosphorylation behavior during beta-cell differentiation. A very exciting find was to identify a key beta-cell transcription factor NKX6.1 among phosphorylated proteins and we first searched for publications with no result, and also no phospho-NKX6.1 antibody is available. We show three serine residues of NKX6.1 (S167, S317 and S334) as targets for a robust phosphorylation during the PP1 to PP2 and even larger one during the PP2 to EN transitions (Fig.5B, lower panel).

To compare the phosphorylation trajectories of Chromogranin A and NKX6.1 we plotted log-values of phosphorylated peptide abundances for both proteins and show that they follow a similar path (Fig.5C). We hypothesize that both phosphorylations either serve as markers of emerging endocrine populations or connected via yet identified signaling network.

Finally, we aimed to show and compare the increase of CHGA and NKX6.1 phosphorylation at the individual amino acid levels and normalized to a change in protein expression (Fig.5D-E). For NKX6.1, S317 or S334 residues phosphorylation changes were detected in all three samples (Fig.5D, blue with error bars), while S167 was detected only in one dataset (Fig.5D, orange). The S317/S334 phosphorylation peaks at PP2 level, while S167 continues to increase gradually till EN stage. Our interpretation of this phenomena, that each phosphorylation of NKX6.1 may represent a separate signaling event and we hope that it will be clarified in future studies. Chromogranin phosphorylation was gradually increasing as cells were transitioning between protocol stages, however a large uptake in PO4 CHGA levels was observed toward the EN endpoint (Fig.5E). We do not know the biological mechanism involved in this post-translational process for Chromogranin A and hope that it will be explored in future studies.

### Validation of proteome and phosphoproteome targets with single-cell RNA sequencing data analysis

An interesting opportunity presented itself when we applied our visualization and t-SNE methods to published single-cell RNA sequencing datasets (14,23), allowing us to validate and place protein IDs to a specific cell sub-population of the differentiating cluster at EN stage (stage 6) (with Veres et al. dataset) or to a timepoint at the differentiation protocol (with Sharon et. al dataset). We created t-SNE plots for both (Fig.6A) and color-coded either differentiation time-point or cell sub-population in them. Upon creating the web-based application, we plotted any protein ID of interest that were selected from those shown in Fig.3B and 3C, and Fig,4D. The selection of protein IDs was to authors’ discretion in order to emphasize our findings and can be expanded by any future researcher. We show (Fig.6B) that stage and lineage-specific markers such as PDX, NKX6-1, GCG, INS, CHGA and SST appear at the appropriate stage and cell populations, confirming the applicability of the webapplication code. Then, we turned our attention to protein IDs that were identified with unsupervised machine learning in Fig.3C. Here (Fig.6C), we show those IDs that appeared on t-SNE plots within a specific stage or cell subpopulation. While few of the IDs were expected based on literature search, we propose that Carboxypeptidase E (CPE,(71), MAN1A (Mannosidase Alpha Class 1A Member 1,(72), and ALDH1A1 (aldehyde dehydrogenase 1 family member A1, (73) deserve a further consideration for the modulation of differentiation protocol efficiency and developmental research. All other protein IDs from Fig.3C were not clearly identified or were at minimal detection levels at both t-SNE plots.

**Figure. 6.**
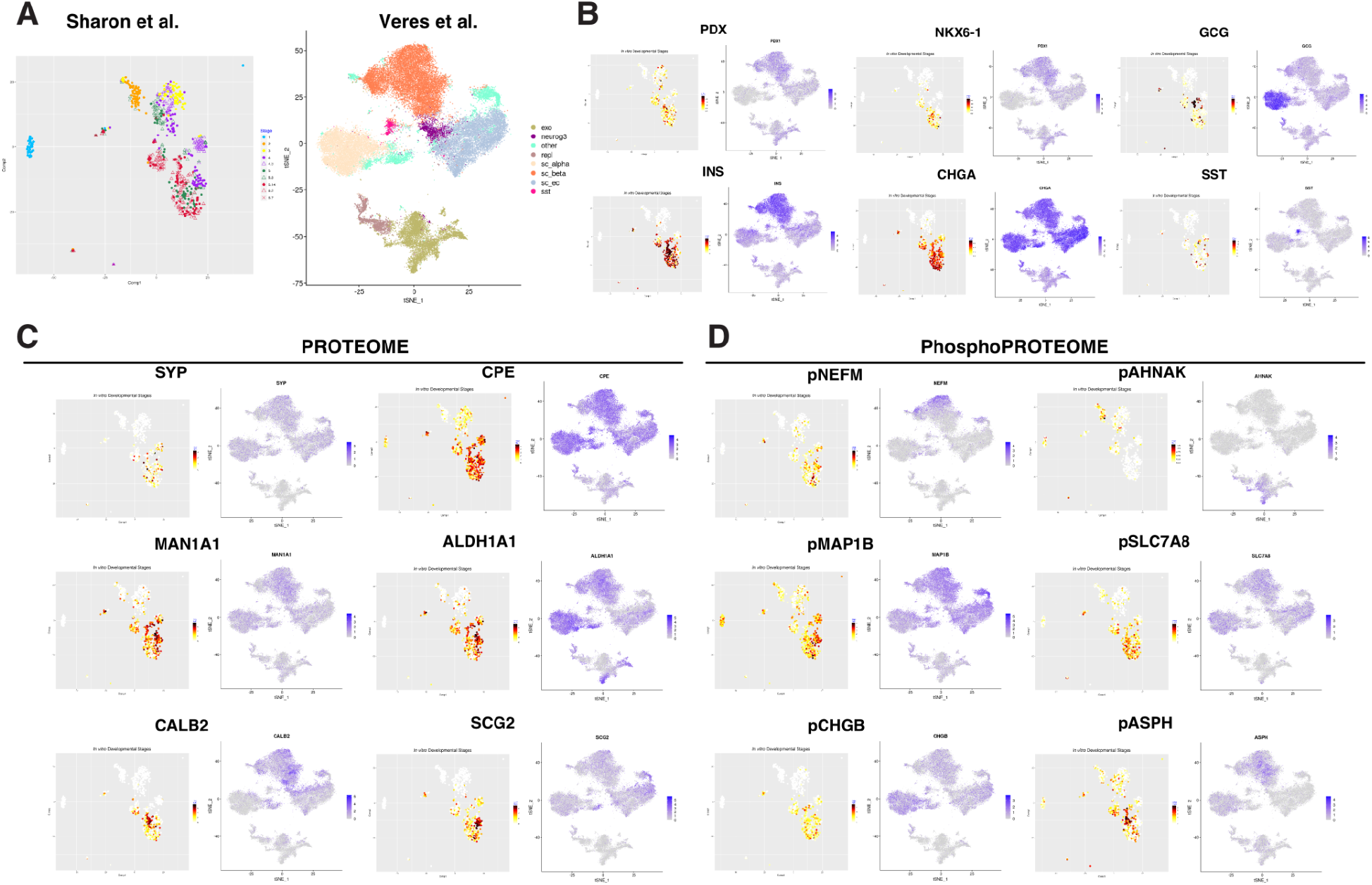
Confirmation of protein and phosphoprotein targets with single-cell RNA sequencing data processing. **A.** Single-cell data from Sharon et al. (23) was analyzed as described in Methods with Shiny R package and visualized with t-distributed stochastic neighbor embedding (t-SNE) plots. Data points represent differentiation protocol stages as follows: 1 is the ESC stage; 2-DE; 3-PGT; 4-PP1; 4.3-day 3 of PP1, 5-PP2, 5.3-day 5 of PP2; 6-EN, 6.2,6.7 and 6.14-days 2, 7 and 14 into the EN stage (maturation stage of the protocol). **B.** Single-cell data from Veres et al. (14) was analyzed as described in Methods with Shiny R package and visualized with t-SNE plots. EN stage (stage 6) cell populations are abbreviated as follows: EXO-exocrine; NEUROG3-Neurogenin 3 (NGN3) progenitors; OTHER-other cells types; REPL-replicating cells/differentiating; SC-alpha-a cells; SC-beta – b cells; SC-EC-enterochromaffin cells; SST-d (delta) cells. **C.** Beta-cell differentiation stage markers expression patterns as projected to the single-cell data t-SNE plots from Figure 6A. **D** Selected proteins from Figure 3C that were identified with unsupervised machine learning as plotted with Sharon and Veres dataset analyzes. **E.** Selected phosphoproteins from Figure 4D for stage transitions PP1 to PP2 and PP2 to EN as plotted with Sharon and Veres dataset analyzes.

Next, we applied the same visualization method to phosphoprotein IDs from Fig.4D. We show that most of the highest-phosphorylated IDs can be associated with the appropriate stage and cell populations, however still there were many that we couldn’t plot either due to low transcript levels or no association with a defined cell population or protocol stage. Phosphorylated protein targets of interest that appeared at later stages of the protocol and within a specific cell population were NEFM (neurofilament medium polypeptide) and MAP1B (Microtubule Associated Protein 1B), both associated with vesicle trafficking in neurons, where NEFM is highly upregulated in a subset of beta-cell population. In exocrine cells, AHNAK-cell cycle mediating protein (74) is highly expressed and was shown to modulate TGFb/Smad signaling pathway, which presents it as an interesting target for a future research in beta-cell development. A highly-expressed SLC7A8 protein (75) (L-type amino acid transporter 2, LAT2) is also highly-phosphorylated at the EN stage, modulates mTOR signaling (76) and associated with islet cells lineage, but we do not find any publication with regard to its phosphorylation in islet development. Same result was for ASPH protein (aspartate β-hydroxylase), where we found several studies for its role in pancreatic cancer but none in pancreas development.

Lastly, we turned our attention to mapping ECM (collagens, laminins), metalloproteases and integrin proteins (Supp. Fig.7). We screened all protein IDs from Fig.3B and Supp. Fig.6 and show only those that were clearly identified on t-SNE plots (Supp. Fig.8). As expected from previous studies we clearly show integrin a-1(ITGA1/CD49a,(14,77) and integrin b-1 (ITGB1/CD29, (78). However, the rest of integrins from Fig.3B were not as clearly detected with the single-cell t-SNE analysis. Similar results were obtained for metalloprotease, laminins and collagen isoforms. We show only those t-SNE plots and protein IDs, where a clear stage or cell population was confirmed. The most prominent stage and cell association was observed for collagen V isoform (COL5A2), where it appears to begin an increased expression at DE stage of the differentiation (Supp. Fig.6), but also is upregulated toward later stages, as shown in t-SNE plots. A recent publication (79) suggested its importance for an increased functionality of stem cell-derived beta and alpha cells in vitro and we are proposing here that collagen V role may be associated with enterochromaffin cell population which is known to positively regulate insulin secretion in pancreas (80,81). It presents an attractive translational target, where collagen V combined with other ECM proteins and biopolymers may improve the maturation and functionality of stemcell derived replacement islets for T1D patients.

## Conclusion

Here we present the analysis of phenome (transcript and protein landscape) dynamics during beta-cell differentiation in vitro from embryonic stem cells, as cell transition through developmental stages in tissue culture. Understanding the change in gene expression at mRNA and protein levels with the addition of phosphorylation PTM data, with the deep resolution of high-throughput sequencing and mass-spectrometry methods was supported by modeling tools and machine learning analysis, implemented in various analytical scenarios here. This work provides researchers with a source of signaling networks in beta-cell differentiation, which can also be applied for the fundamental understanding of pancreatic development and the improvement of cell differentiation processes.

We show that amplitude and the number of changes in gene expression at each stage of the protocol not always directly correlate between mRNA and protein levels. Phenome exhibits various dynamic temporal profiles as cells progress through the protocol, underlining the differences between protein abundances and changes in mRNA levels. Our datasets generated with the most recent protein mass spectrometry and mRNA sequencing methods are providing comparable numbers of unique genetic IDs, allowing researchers to focus on the analysis of network and gene expression correlations with a high level of precision. Thus, the inherent complexity of modeling signaling networks within a dynamic system of stem cell differentiation attributes to biological differences and less so to the methodological errors or their resolution. Increasing the depth of both the proteome and the transcriptome in this study allows identification with significant levels of confidence of protein markers associated with a specific stage of differentiation, which may previously have been missed. We compared our findings of main protein stage markers (PDX1, NKX6.1, INS and others), and confirmed the presence of signaling pathways already utilized by beta-cell differentiation protocol in vitro while using these markers as anchors in building our networks and identification of new targets within.

Our interpretation of mRNA/Protein dynamics and profiles leads to the conclusion that the study of cell population dynamics in culture requires phenomic analysis for each gene of interest. This notion entails researchers be mindful of the cell population behavior as a whole, which is equally vital in highly dynamic systems when compared to a single cell decision making and differentiation (82). Our findings of the correlation coefficient for proteome vs. transcriptome are within the reported range of values 0.4-0.6 as previously reported (83–86).

The funnel profile of mRNA expression of ESCs during the protocol, with the total number of the altering transcripts, shows that stem cells enter the differentiation process while exhibiting a significant transcriptional noise, and towards the end of the process cells may remain in a balanced mode between stochastic and deterministic states of fate determination (82,87,88). This observation is emphasized by the point that until the entry into PP1 to PP2 transition cell population remains rather homogenous (SOX17+ cells approximately 80% and PDX-1-positive cells 90% of the sample). Once we establish that cells remain in a fluid state concerning their fate decision still in the middle of the differentiation protocol, each perturbation of the PDX-1-positive population carries more significance for the emergence of endocrine cells. Different cell populations at the EN stage can attribute to the transcriptional noise, but at the PP1 stage, this noise indicates that cells haven’t fully being committed to the endocrine lineage. It brings more importance to finding new protein targets to be perturbed for better endocrine lineage determination of the PDX-1-positive cells or cells at the DE, PGT, or even the PP2 stages of the protocol.

Primary motivation behind this project was to identify new signaling pathways involved in beta-cell differentiation and developmental fate determination. We conclude that at the early stages (DE, PGT, PP1, and PP2) of the protocol differentially expressed proteins represent distinct biological processes that may or may not be related, whereas at the EN stage their correlation may indicate a cooperative function in endocrine differentiation. These observed patterns of mRNA and protein dynamics of a specific differentially expressed gene can be attributed to cell type specialization and establishment and serves as yet another indication of an initial stochastic behavior of cell population that persists until the PDX-1-positive or even NKX6.1-positive cells. This pattern reverses upon the emergence of CHGA expression, which indicates the setting of endocrine lineage fate in culture. It also points to a complex cellular attractor landscape (82), where at the PP1 stage (PDX1-positive cell population) more than one signaling pathway leads to endocrine lineage. New proteins, identified with machine learning approaches through anchor protein clustering and profile similarity, present an opportunity for the identification of new signaling pathways involved in stem cell fate determination and betacell development.

We show that single-cell RNA sequencing data and proteomics do not contradict each other and in general, support each other conclusions. Having said so, our proteoform datasets presented many protein and phosphoprotein targets that could have been missed if studied only at the transcriptome level. For example, integrin interaction with ECM are fundamental for islet formation and functionality (89), and we show many ECM components to be associated with a specific lineage marker at the proteome level, but not clearly mapped on the single-cell transcriptome landscape.

Our findings and resource data here can be utilized in furthering single cell studies of genetic networks and critical regulators that govern endocrine differentiation. These studies (90–93) were aimed on defining cell populations within fetal and adult islets, finding cell-specific protein and transcript markers for a specific cell within the islet and deciphering the spatiotemporal signaling of their development. Combining single cell approaches for transcriptomics sample processing and analysis (10,94), in-depth proteomic and phosphoproteomic studies (95), with single cell proteomics (96) methods can generate new targets for beta cell differentiation protocols.

Our discovery on previously unknown phosphorylations of Chromogranin A and NKX6.1 open the door for future studies of translational networks, governed by NKX6.1 (97,98), and endocrine secretion machinery where Chromogranin A plays an important role in peptide hormones family and islet cells function (97,99). While we point to NKX6.1 phosphorylation as suggested target for understanding molecular signaling which leads to the emergence of beta and other islet cells, we do not know a significance for the rapid increase in Chromogranin A phosphorylation at late stages of differentiation protocol.

We present our datasets here for the further study of developmental biologists and stem cell translational researchers for differentiation protocol betterment and specialization for a cell type desired.

## Experimental procedures

### Beta-cell differentiation protocol in vitro

HUES8 human embryonic stem cells (RRID:CVCL_B207), included in NIH human embryonic stem cell registry (registration number 0021), were differentiated into beta-like cells in suspension as described previously (100). Briefly, 150×10^6^ HUES8 ESC cells suspended in 300 ml of supplemented mTeSR1 (StemCell Technologies) and grown in clusters in spinning flasks (70rpm, 37°C, 5%CO_2_, Corning) for 48 hours prior to the beginning of a stepwise differentiation protocol:

ESC to DE transition – 24 hours in S1 medium supplemented with Activin A (100ng/ml, R&D Systems) and CHIR99021 compound (14μg/ml, Stemgent), followed by 48 hours without CHIR99021.

DE to PGT transition– 72 hours in S2 medium supplemented with KGF (Peprotech, 50ng/ml).

PGT to PP1 transition – 48 hours in S3 medium supplemented with KGF (50ng/ml), LDN193189 (Sigma, 200nM), Sant1 (Sigma, 0.25μM), retinoic acid (Sigma, 2μM), PDBU (EMD Millipore, 500nM) and ROCK inhibitor Y-27632 (Tocris, 10μM).

PP1 to PP2 transition – 5 days in S3 medium supplemented with KGF (50ng/ml), Sant1 (0.25μM), retinoic acid (RA, 0.1μM), Y-27632 (10μM) and Activin A (5ng/ml).

PP2 to EN transition– 7 days in BE5 medium supplemented with Betacellulin (Thermo Fisher Scientific, 20ng/ml). XXI (DNSK, 1μM), Alk5i (Axxora, 10 μM) and T3 (DNSK, 1μM). Sant1 (0.25μM) were added in the first three days, and retinoic acid was added at 0.1μM in the first three days, then at 0.025μM.

Whole cellular clusters (30 x10^6^ cells for proteomics and 1 x10^6^ cells for RNA sequencing) were collected and washed with pre-chilled PBS (-Ca^2+^/-Mg^2+^, Corning), snap frozen (−80°C) for proteomics analysis or dissolved in 300 ul of RNAlater reagent (Qiagen).

### Flow Cytometry

Differentiated cell clusters or islets were dispersed into single-cell suspension by incubation in TrypLE Express at 37°C, fixed with 4%PFA on ice for 30 min, washed once in PBS, and incubated in blocking buffer (PBS+0.1% Triton X-100+5% donkey serum) on ice for 1 hr. Cells were then re-suspended in blocking buffer with primary antibodies and incubated at 25°C for 90 min. Cells were washed twice in blocking buffer and were then incubated in blocking buffer with secondary antibodies on ice for 2 hr. Cells were then washed thrice and analyzed using the LSR-II flow cytometer (BD Biosciences). Analysis of the results was performed using FlowJo software. Chromogranin A (101,102) levels above the threshold of 35% of co-expression with NKX6.1 within clusters (NKX6.1+/CHGA+ %) were used as a reference checkpoint during the differentiation, for internal Go/No-Go decision making in beta-cell production.

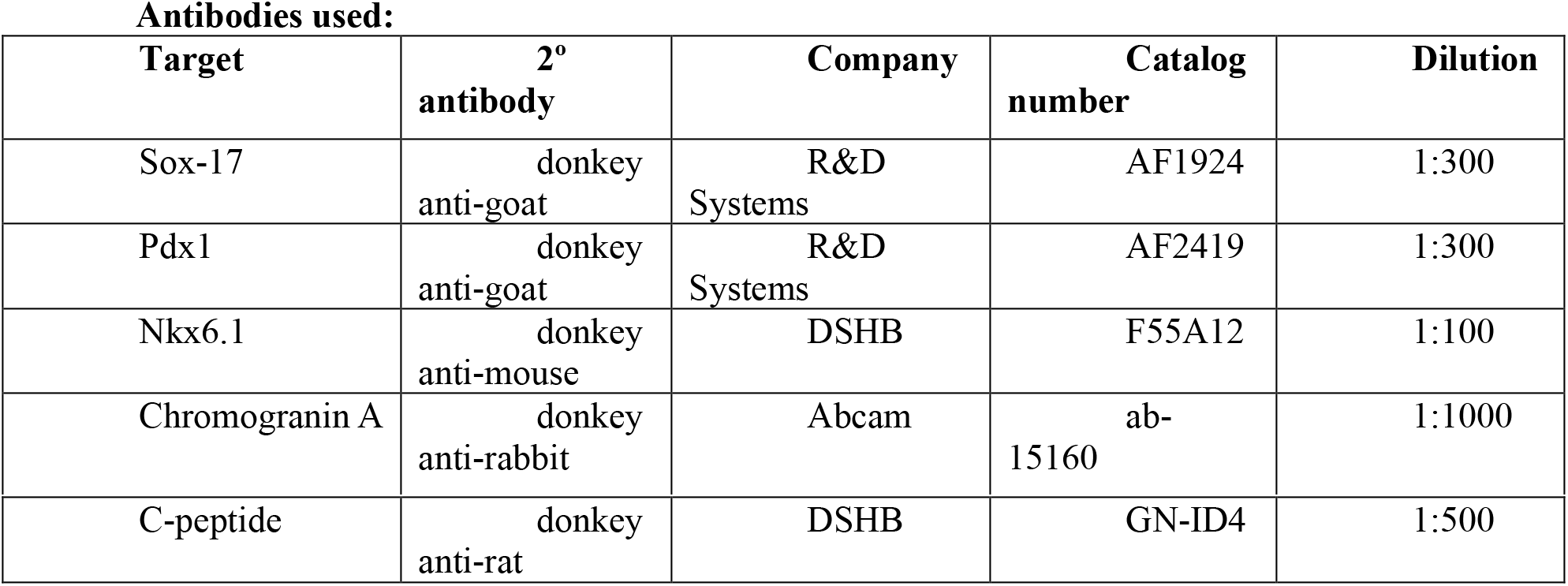

Antibodies used:
| Target | 2° antibody | Company | Catalog number | Dilution |
| --- | --- | --- | --- | --- |
| Sox-17 | donkey anti-goat | R&D Systems | AF1924 | 1:300 |
| Pdx1 | donkey anti-goat | R&D Systems | AF2419 | 1:300 |
| Nkx6.1 | donkey anti-mouse | DSHB | F55A12 | 1:100 |
| Chromogranin A | donkey anti-rabbit | Abcam | ab-15160 | 1:1000 |
| C-peptide | donkey<br>anti-rat | DSHB | GN-ID4 | 1:500 |

### Proteomics and sequencing sample data acquisition

#### Sample lysis, DNA shearing, digestion and TMT 6-plex labeling

Cell clusters samples from each of six differentiation steps were submitted for proteomics analysis in 50ml conical plastic tubes, after brief wash with PBS and snap freeze in −80°C. Each was brought to 5% SDS by volume (~1 ml total final volume) and subjected to complete lysis and DNA shearing using a Covaris Model S220 Focused Acoustic device (Covaris, Woburn MA) in 1ml glass Covaris vials. The samples were reduced (TCEP added to 20mM, 90°C for 20 minutes), cooled to room temperature (RT), and alkylated (Iodoacetamide added to 50mM, room temperature in the dark for 30 minutes). The individual samples were subjected to detergent removal and enzymatic digestion using the S-Trap (Protifi, Huntingtion, NY) affording the workflow facile removal of SDS and fast efficient digestion time (~1 hour) with clean peptides eluted directly from the device.

Each of the six digested samples were dried to near full dryness on a SpeedVac (Eppendorf, Germany) and resuspended in 100ul 50mM TEAB. Following TMT six-plex Isobaric Label reagent set directions (ThermoFisher Scientific, Carlsbad, CA) each sample was labeled, allowed to react for 1 hour at RT, the reaction was quenched with the addition of 5% hydroxylamine, and the six TMT labeled samples were then combined. This final mixture was dried fully in the Speedvac for subsequent processing.

### ERLIC separation and Mass spectrometry analysis

After trypsin digestion and TMT labelling, the combined peptides were separated on an Agilent 1200 HPLC system (Santa Clara, CA) using PolyWAX LP column (PolyLC, Columbis MD) 200×2.1 mm, 5um, 300A running under ERLIC mode conditions (Electrostatic Repulsion Hydrophilic Interaction Chromatography). Peptides were separated across 90 min gradient from 0% buffer A (90% Acetonitrile 0.1% Acetic Acid) to 75 % buffer B (30% Acetonitrile, 0.1% Formic Acid) with 20 fractions collected by time. Each fraction was dried in SpeedVac (Eppendorf, Germany) and resuspended in 0.1% formic acid solution before analysis by mass spectrometry.

Each ERLIC fraction was submitted for a single LC-MS/MS experiment that was performed on a Tribrid Lumos Orbitrap (Thermo Fischer Scientific, Carlsbad, CA) equipped with EASY nLC-1000 UHPLC pump. Peptides were concentrated and desalted on a 5 cm long by 150 μm inner diameter microcapillary trapping column packed with Reprosil-Pur 120 C18-AQ media (5 μm, 120 Å, Dr. Maisch GmbH, Germany) then resolved and eluted from the analytical column of ~25 cm of Reprosil-Pur 120 C18-AQ media (1.9 μm, 120 Å, Dr. Maisch GmbH, Germany). These columns were fabricated in-house from glass capillary tubing that had KASIL frits (potassium-silica) baked in place. These were cut to length and the glass capillary ends polished using the Capillary Polishing Station (the CPS) (ESI Source Solutions, Woburn, MA), the conformity of these ends confirmed using a USB microscopy station, with the capillary tubing then flushed clean with ethanol flow and sonication. Once constructed the capillary columns were packed with reversed phase media using a pressure vessel and nitrogen gas. Wherever possible, PEEK Cheminert fittings (VICI Instruments) were used to ensure optimal chromatographic performance. Separation was achieved with a gradient from 5–27% ACN in 0.1% formic acid over 90 min at 150 nl/min. Electrospray ionization was enabled through applying a voltage of 2 kV using a custom nanospray tip assembly (ESI Source Solutions) which held the steel micro union for the high voltage liquid junction at the end of the analytical column and sprayed from uncoated fused silica Picotips (New Objective, MA) with dimensions of 360 μm OD, 20μm ID and 10μm spray tip. Background ions were reduced and signal to noise ratio was enhanced at the instrument inlet by using an Active Background Ion Reduction Device (ABIRD) (ESI Source Solutions, Woburn, MA). The Tribrid Lumos Orbitrap was operated in data-dependent mode for the mass spectrometry methods. The mass spectrometry survey scan was performed in the Orbitrap in the range of 400 –1,800 m/z at a resolution of 6 × 10^4^. The TOP20 ions were subjected to HCD MS2 event in Orbitrap part of the instrument. The fragment ion isolation width was set to 0.7 m/z, AGC was set to 50,000, the maximum ion time was 150 ms, normalized collision energy was set to 37V and an activation time of 1 ms for each HCD MS2 scan with resolution of 5×10^4^ as well as to CID MS2 scan with collisional energy at 35V and Isolation window of 1.6 Da.

### Mass spectrometry analysis

Raw data were submitted for analysis in Proteome Discoverer 2.2.0388 (Thermo Scientific) software. Assignments of MS/MS spectra were performed using the Sequest HT algorithm by searching the data against the protein sequence database including all entries from the Human Uniprot database (Human SwissProt 16,792 2016 and Human Uniprot Trembl 2016) and other known contaminants such as human keratins and common lab contaminants. Sequest HT searches were performed using a 10-ppm precursor ion tolerance and requiring each peptides N-/C termini to adhere with Trypsin protease specificity, while allowing up to two missed cleavages. 10-plex TMT tags on peptide N-termini and lysine residues (+229.162932 Da) was set as static modifications while methionine oxidation (+15.99492 Da). A MS2 spectra assignment false discovery rate (FDR) of 1% on protein level was achieved by applying the targetdecoy database search. Filtering was performed using a Percolator 64bit version (103). For quantification, a 0.02 m/z window centered on the theoretical m/z value of each the six reporter ions and the intensity of the signal closest to the theoretical TMT m/z value was recorded. Reporter ion intensities were exported in result file of Proteome Discoverer 2.2 search engine (ThermoFisher Scientific, Carlsbad, CA) as an excel tables, after filtering to a maximum of 40% of co-isolation for the parent ion, all quantitation above that threshold were removed from further analysis for quantitation purposes. The total signal intensity across all peptides quantified was summed for each TMT channel, and all intensity values were adjusted to account for potentially uneven TMT labeling and/or sample handling variance for each labeled channel.

### Data Processing

We pooled all identified proteins from all samples of the proteomics data to determine the protein coverage, which resulted in 12943 unique protein ID identified. This set had 742 proteins with missing abundance values which were removed from the dataset resulting in a full complement of 12201 proteins for further processing and analysis. Statistical analysis was performed on the subset of proteins that are present at all stages reducing the original number of proteins to 6785. Protein abundances of all samples were normalized using the quantile method and transformed with a log base 10 function.

Raw RNAseq data reads were aligned to the human genome using the RSEM algorithm (104) and the expected gene counts were used for further processing. These were converted to counts per million and only genes with approximately 10 counts (depending on library size) in five or more libraries were retained. Counts across samples were normalized by calculating scaling factors based on trimmed mean of M-values (TMM) with library functions defined in the edgeR package (105).

### Determination of Differentially Expressed Proteins and Genes

The statistical method used to test for differential expression of the proteomics data was as described in (106) and implemented in the limma library (107) of the Bioconductor project (108). The method is based on fitting a linear model to each of the proteins in the dataset and uses a moderated t-statistic to increase the statistical power for small sample sizes. The p-values of differences in expression for each protein were adjusted to control for small p-values that can arise by chance when many tests are performed (multiple testing adjustment). The adjusted p-values are Benjamini-Hochberg q-values (109). The genomics data was preprocessed with the *VOOM* function (110) from the LIMMA package (106) and statistically analyzed with the linear model method from LIMMA as described for the proteomics data.

### Annotation of Proteins

Proteins were annotated with gene symbols and protein names using information contained in the UniProt KB database and the Gene database from NCBI (https://www.ncbi.nlm.nih.gov/). A Python script was used to query UniProt KB with the accession numbers of the proteins identified in the proteomics experiments. The Gene database from NCBI was queried by R methods for gene aliases (111). The collected annotation information was used in a lookup table in R to interconvert between protein UniProt accession ID, gene symbol, and protein name.

### Correlation between Proteins and Genes for each Development Stage

A protein abundance matrix (P) and a gene expression matrix (R) were prepared where gene accessions mapped one-to-one on protein accessions resulting in matrices with 8479 rows. Many-to-one or one-to-many mappings were aggregated by taking the mean over multiplicative expressions or abundances. Pearson correlations were calculated with the *COR* R function between the 6 matrix columns, representing the 6 development stages. The correlation values were visualized in a heat-map by mapping each correlation value to a color in the range of blue, white, and red.

### Correlation of DE Proteins and Corresponding Gene transcripts

The abundances of proteins differentially expressed at the 95% level between sequential development stages were used to determine the correlation with the expression of the corresponding genes. There were 108 differentially expressed proteins from all binary comparisons between consecutive stages. Correlations were determined separately for the subset of differentially expressed proteins between consecutive stages. The resulting correlation matrices were visualized with a heatmap.

### Scatterplots of Proteins and Genes

Abundances and Expressions in matrices P and R, respectively, were plotted in a series of scatterplots, one for each of the 6 assayed stages of beta-cell development from HUES cells. The scatterplots were overlaid with lines of identical densities of abundance-expression tuples. Differentially expressed proteins were marked as red dots and label with their symbols.

### Principal Component Analysis of Protein Abundance and Gene Expression

The principal component analysis on protein abundance and gene expression matrices was carried out using the *PRCOMP* R function with scaling. The matrix of variable loadings, the stage variables expressed as linear combinations of base vectors in the direction of maximal variance, was plotted as a line graph, with a profile line for each stage variable.

### Sub-clustering and profile plotting

Protein abundance profiles over the beta cell development stages were clustered using the affinity propagation methods of unsupervised machine learning (112,113). The algorithm found 68 clusters, which were searched for proteins of interest. The cluster with the selected protein was sub-clustered using the *k*-means algorithm. Optimal cluster numbers for the *k*-means were determined by trial and error to find close-fitting sub-clusters with similar abundance profiles across the stages to the selected stage marker profile.

### ENRCHR analysis of protein clusters

Proteins of interest were clustered by individual criteria of the study (by beta-cell protocol stage, stage-specific markers or by machine learning). Protein ID were imported into ENRCHR (http://amp.pharm.mssm.edu/Enrichr/) online search tool (114,115). ENRCHR platform allows multiple choice selection of analytical outputs and for the purpose of the current study, transcription protein to protein (PPIs) interactomes were selected, for the identification of possible signaling networks coordination with proteins of interest.

### Analysis and visualization of single-cell RNA sequencing datasets

Visualization and querying of single cell (sc) RNAseq data for Sharon, N. et al., (23). The RNAseq data were normalized and clustered with SIMLR as described above, and the Shiny R package (https://shiny.rstudio.com/) was used to render querying and plotting functions results in a dynamic web application by binding web instructions to R functions acting on the data object. Visualization and querying of sc-RNAseq data for Veres, A. et al., (14) was done with Shiny R package as well. The normalized and clustered data from both datasets were downloaded from the GEO repository and the UMAP coordinates (116) extracted. These were used to plot the pancreatic islet cell clusters and highlight gene ID expressions across different lineages of cells at EN stage (stage 6 at Sharon data) with the Veres dataset or at different stages of the differentiation protocol (Sharon dataset).

### Coefficient of Variation

The coefficient of variation for protein abundance and gene expression was calculated for each replicate at each stage of the developmental program and the resulting values were displayed as bar plots.

### Funnel Plot

The differentially expressed proteins of each of the binary comparisons between consecutive development stages were separated into upregulated and downregulated proteins/genes. These were plotted as positive or negative log fold change values categorized with stage number. Smoothed curves were drawn separately through the maximal value points of positive and negative Log (10) fold changes.

### Software packages

Workflows for processing of proteomics and genomics data, statistical analysis of such data and the visualization of the results of such analysis were developed in scripting programs written for the R statistical programming environment (108,111). Use was specifically made of the MSnbase library to provide a convenient data structure for proteomics data (117). Functions from the LIMMA R package(https://bioconductor.org/packages/release/bioc/html/limma.html) were used for statistical modeling for both the proteomics and genomics data. In the case of the genomics data the *VOOM* R function, contained in the edgeR package (https://bioconductor.org/packages/release/bioc/html/edgeR.html), was used to preprocess the count date before the linear modeling method was used.

## Acknowledgments

This work was supported by and AstraZeneca pharmaceutical company collaboration with Harvard Stem Cell Institute (HSCI) and Melton group. We express our sincere gratitude to Dr. Douglas A. Melton and members Melton laboratory at HSCI in Cambridge, MA for their dedication and support for this project. We thank Renee Robinson, Elise Engquist, Chunhui Xie, Yu Yi, Kaleigh Biles and Benjamin Rosenthal for their help and hard work collecting and preparing samples. We acknowledge a dedicated work of Dr. Claire Readron and Harvard FAS sequencing group. We thank Dr. Michele Clamp, Dr. Nadav Sharon, Dr. Bjorn Tyberg and Astrozeneca bioinformatics group for the help discussing our approach and providing with manuscript comments. We also thank Nils Bergenheim from Astrozeneca for the help managing research collaboration. We thank Nathaniel Jorgenson, Cellaria internship student, for data analysis. D.S. is currently employed at Cellaria Inc.

## Data availability statement

The data that support the findings of this study are openly available in PRIDE Archive at https://www.ebi.ac.uk/pride/, reference number #PXD021521 and in MassIVE database https://massive.ucsd.edu/, with DOI: 10.25345/C5PJ20.

